# Assembly and phylogeographic analysis of novel *Taenia solium* mitochondrial genomes reveal further differentiation between and within Asian and African-American genotypes

**DOI:** 10.1101/2022.03.10.483888

**Authors:** Gabriel Jiménez-Avalos, Alina Soto Obando, Maria Solis, Robert H Gilman, Vitaliano Cama, Armando E Gonzalez, Hector H García, Patricia Sheen, David Requena, Mirko Zimic, the Cysticercosis Working Group in Peru

**Affiliations:** Laboratorio de Bioinformática, Biología Molecular y Desarrollos Tecnológicos. Laboratorios, de Investigación y Desarrollo., Facultad de Ciencias y Filosofía., Universidad Peruana Cayetano Heredia., Av. Honorio Delgado 430., San Martín de Porres, 15102., Lima, Perú; Department of International Health., Bloomberg School of Public Health., Johns Hopkins University., 615 North Wolfe St., Room 5515., Baltimore, 21205., Maryland, USA; Division of Parasitic Diseases and Malaria., Center for Global Health., Centers for Disease Control and Prevention., 1600 Clifton Rd. MS D-65., Atlanta, GA 30329. The USA; Facultad de Medicina Veterinaria., Universidad Nacional Mayor de San Marcos., Av Circunvalación 28, San Borja, 15021, Lima, Peru; Departamento de Microbiología., Universidad Peruana Cayetano Heredia, Av. Honorio Delgado 430., San Martín de Porres, 15102., Lima, Perú; Cysticercosis Unit, Instituto Nacional de Ciencias Neurológicas, Jr. Ancash 1271., Cercado de Lima 15003., Lima, Perú; Laboratory of Cellular Biophysics., The Rockefeller University., 1230 York Avenue., New York, NY 10065., USA

**Keywords:** Phylogenetics, Phylogeography, Haplotypes, Taeniasis, Cysticercosis, Genetics, Genomics, Genetic Epidemiology, Mitochondrial genome

## Abstract

**Background:** *Taenia solium* is a parasite that hampers human health, causing taeniasis and cysticercosis. The genetic variability in its mitochondrial genome is related to the geographical origin of the specimen. Two main genotypes have been identified: The Asian and the African-American. The geographic genetic variability is expected to cause different clinical manifestations. Thus, characterizing differences between and within genotypes is crucial for completing the epidemiology of *T. solium* diseases.

**Methods/Principal Findings:** Here, two Peruvian (one complete and one partial; 7,811X and 42X of coverage, respectively) and one Mexican (complete, 3,395X) *T. solium* mitochondrial genomes were assembled using the Chinese reference. Variant calling with respect to the reference was performed. Thirteen SNPs that involved a change in the amino acid physicochemical nature were identified. Those were present in all the assembled genomes and might be linked to differences in aerobic respiration efficiency between Latin American (African-American) and Asian genotypes. Then, phylogeographic studies were conducted using Cytochrome C oxidase subunit I and cytochrome B from these genomes and other isolates. The analysis showed that Indonesian samples are the most ancient and related to the modern *T. solium* ancestor of the Asian genotype. Finally, a consistent subdivision of the African-American genotype into two subgroups was found. One subgroup relates to East African countries, while the other is West Africa. The East African linage suggests a previously unnoticed influence of the Indian Ocean trade in the genetic structure of Latin America *T. solium*.

**Conclusions/Significance:** Overall, this study reports novel mitochondrial genomes valuable for further studies. New Latin American SNPs were identified and suggest metabolic differences between parasites of the Asian and African-American genotypes. Moreover, the phylogeographic analysis revealed differences within each genotype that shed light on *T. solium’s* historical spread. Overall, the results represent an important step in completing *T. solium* genetic epidemiology.

**Author Summary:** *Taenia solium* is a human parasite that causes taeniasis and cysticercosis. Eradicated from developed countries, they are still a public health problem in developing nations. *T. solium* differences in the mitochondrial genetic material depend on its geographical origin. This is expected to cause different clinical manifestations. Despite the importance of genetics to the epidemiology of *T. solium* diseases, few efforts have been made to assemble and compare their genomes. We aimed to help fill this knowledge gap by assembling three mitochondrial genomes from Latin America and comparing them to the Chinese reference. Additionally, two genes from the Latin American genomes and from other isolates were employed to assess *T. solium* genetic distribution. We found thirteen mutations with respect to the Chinese genome present in all Latin American samples, which involved a change in the amino acid physicochemical nature. Those might be causing metabolic differences between Asian and Latin American parasites that could change their affinity to specific human tissues. Moreover, we determined that Indonesian samples are the most ancient and related to the modern *T. solium* ancestor. Finally, we identified a previously unnoticed influence of East African countries in *T. solium* phylogeny, with which our assembled genomes are closely related.

## INTRODUCTION

*Taenia solium* is a parasite responsible for two critical diseases in humans: taeniasis and cysticercosis. The former refers to infection with the adult stage of the parasite. The latter is the infection with its larvae and represents a major risk to human health. Cysticercosis could progress to the central nervous system causing neurocysticercosis, the leading cause of acquired adult epilepsy in developing countries [1]. Humans are the only known definitive host, harboring the adult tapeworm and releasing infectious eggs to the environment [2–4].

*T. solium* has spread globally, being endemic and highly prevalent in Asia, Africa, and Latin America [5]. Interestingly, it has been shown that *T. solium* intraspecies variability in the mitochondrial genome is strongly related to the geographical origin of the specimens [6–9]. Two main genotypes have been identified, the Asian and the African-American [6]. This geographic genetic variability is expected to result in clinical heterogeneity in *T. solium* diseases between regions [10]. Therefore, an exhaustive study of it is crucial for completing the epidemiology of taeniasis and cysticercosis [6,11]. Despite this, there is a lack of characterization of differences between whole *T. solium* mitochondrial genomes from different genotypes due to the few assembled genomes available.

Mitochondrial genes, specially Cytochrome C oxidase subunit 1 (COX1) and cytochrome B (CYTB), have proved to be useful markers to assess intraspecies variability and phylogeography of *T. solium* [6–9,12–14]. Both genes have low variability [6]; however, CYTB is suggested to be slightly more variable than COX1 in *T. solium* [6,15]. The extremely low variability of COX1 limits its use in intraspecies studies of this parasite, being CYTB more suitable for this purpose [15]. Comprehensive analysis of multiple *T. solium* mitochondrial genomes from different origins will confirm if CYTB is the absolute best sequence for this kind of analysis.

The geographic distribution of *T. solium*’s genotypes was shaped by human migrations and trade [6,7,9,12–14,16]. For example, the similarity and gene flow between Latin American, African, and Philippine *T. solium* populations resulted from the European maritime trade routes of the XV and XIX centuries. Furthermore, the sympatric coexistence of the Asian and African-American genotypes in Madagascar is explained by two independent human groups that migrated to this island and introduced both lineages [9,12]. In that sense, geographic genetic differences (or similarities) between *T. solium* populations depend on the connection degree of those places’ human groups. These differences can be used to assess the impact of human migration and trade on the spread of this parasite [9,12,14], which is essential to prevent its dissemination. However, there is also a lack of research on differentiation within each genotype, although previous work has suggested, for instance, that two sublineages could exist within the African-American genotype [8]. Including newly assembled sequences and the ones reported worldwide in phylogeographic studies could help fill this knowledge gap.

Hence, the present study assembled and annotated the *T. solium* mitochondrial genomes of two Peruvian and one Mexican isolate. Those and the Chinese reference mitochondrial genome were compared to provide a detailed characterization of the differences between Asian and African-American genotypes’ genomes. Finally, the COX1 and CYTB sequences from these isolates and from others reported worldwide were included in a phylogeographic reconstruction to analyze differentiation within the Asian and African-American genotypes. COX1 and CYTB were used as they are the genes with the most *T. solium* sequences available from diverse geographical origins.

## MATERIALS AND METHODS

### Assembly and annotation of the mitochondrial genomes

There are currently three Latin American *T. solium* mitochondrial reads available in the NCBI Sequence Read Archive. Two from Peru [17], and one from Mexico (NCBI BioProject: PRJNA170813). Additionally, an assembled reference mitochondrial genome from a Chinese isolate has been published [18].

Whole-genome sequencing reads of the three Latin American samples were collected (Accession codes: SRR644531, SRR650708, and SRR524725). Each sample was independently mapped, using as reference the *T. solium* reference mitochondrial genome of the Chinese isolate (GenBank ID: NC_004022). The mapped reads were subjected to quality control in FastQC v.0.11.9 (http://www.bioinformatics.babraham.ac.uk/projects/fastqc/) and trimmed using a quality threshold of 0.001 in CLC Genomics Workbench v. 21.05.5 (https://digitalinsights.qiagen.com/). These cleaned reads were re-mapped against the reference for each sample. The average coverage of this re-mapping was computed using the pileup script of BBtools v. 38.91 (http://sourceforge.net/projects/bbmap/). The consensus sequences were extracted, inserting the ambiguity symbol “N” to handle conflicts in low coverage regions and inserting the most frequent base (voting) when conflicts occurred in high coverage sections.

In addition, the cleaned reads were *de-novo* assembled to detect genetic rearrangements. We also used CLC Genomics Workbench to perform quality control on each contig (identifying chimeric sequences, misassemblies, and artifacts), measure the depth coverage and %GC content per contig, and calculate the N50 of the assembly.

The consensus sequences of each sample were manually curated to correct misassemblies, which resulted in the final assembled genomes being used in further steps. The annotation of the final assembled genomes was performed using CLC Main Workbench v.21.05.5 using the annotation of the Chinese mitochondrial genome as a reference.

### Variability and selective pressure

Variant calling of the assembled Peruvian and Mexican mitochondrial genomes was performed by whole-genome multiple alignment in the software Mauve v. 2.4.0 [19], using the Chinese *T. solium* mitochondrial genome as reference and default parameters. Single Nucleotide Polymorphisms (SNPs) were manually curated to rule out possible sequencing, alignment, or variant calling errors; and classified into three categories: synonym, non-synonym, and mutations in non-coding regions. This was performed using the software DNAsp v. 6.12.03 [20]. SNPs were graphically represented on a scaled circular map using Circos v. 0.69 [21].

The variability of the 12 protein-coding genes was evaluated as the level of sequence difference (D) [22] for each pairwise combination of the genomes from China, Puno, Huancayo, and Mexico. Each of these genes was re-aligned using the web implementation of the EMBOSS Needle algorithm [23]. D was computed as D = 1 -(M/L), where M is the number of invariant sites and L is the difference between the alignment length and ambiguous bases.

The Ka/Ks ratio of each protein-coding gene against the Chinese reference sequence was computed to evaluate the level of selection pressure and evolutionary adaptation of the *T. solium* mitochondrial genomes. For this purpose, the multiple genome alignment was split into 12 protein-coding gene alignments and submitted to the DNSsp software to calculate the ratio.

### Phylogenetic analysis

COX1 and CYTB mitochondrial genes from the genomes assembled and from different complete gene sequences reported worldwide were employed to perform a phylogenetic reconstruction. This comprises a total of 45 *T. solium* COX1 and 31 CYTB sequences available in GenBank, including, as the outgroup, sequences from *T. saginata* (COX1: AB066495.1 and NC_009938.1; CYTB: AB066581.1 and NC_00938. 1), *T. asiatica* (COX1: AB066494.1 and NC_004826.2; CYTB: AB066580.1 and NC_004826.2) and *Echinococcus multilocularis* (COX1 and CYTB: NP_000928.2).

Multiple global alignments were performed independently for COX1 and CYTB in the software MAFFT v. 7 [24,25], using the progressive G-INS-1 method. The informative coding regions of the alignments were extracted using the Gblocks server v. 0.91 [26,27] with default options.

Phylogenetic analysis was conducted for COX1 and CYTB, separately, using the Maximum Likelihood (ML) and Bayesian Inference (BI) algorithms. For ML, RaxML v. 8.2.12 [28] was used with the GTRCAT model, and 1000 bootstrap replicates to estimate the robustness of the branches. BI was conducted in BEAST2 v. 2.6.1 implemented on the CRIPRES online server platform [29].

The evolutionary model was estimated with jModelTest2 v. 2.1.6 for COX1 (GTR+G with four gamma categories) and CYTB (GTR+I, I = 0.5720) [30], using the Akaike criterion correction. A basic coalescent model for demographic history and a relaxed molecular clock model (uncorrelated lognormal) during the Markov Chain Monte Carlo (MCMC) process were employed. The substitution rates were set to 0.0225 and 0.0195 substitutions per site per million years for COX1 and CYTB, respectively, according to the genetic distance computed for *T. saginata* and *T. asiatica*, which diverged 1.245 [0.78, 1.71] million years ago (Myra) [9,16]. The analysis was run for 50 million generations, sampling every 5000 states and using a burn-in of 10% to obtain an effective sample size (ESS) greater than 200. Lastly, a Maximum Clade Credibility tree (MCC) in TreeAnotator v. 2.6.0 [31] was generated.

### Haplotype network

Haplotypes were identified in DnaSP using as input the multiple alignments of COX1 and CYTB previously generated and considering the total number of mutations as nucleotide substitutions. To prevent adding ambiguity to the network, ambiguous nucleotides were not considered gaps. The haplotype networks were calculated by Median Joining using the software Networks v.10 (fluxus-engineering.com) [32]. The genetic differentiation (ϕst) between the African-American subclades 1 and 2 was calculated using a haplotype distance matrix in Arlequin v. 3.5.2.2 [33].

## RESULTS

### Genome assembly and annotation

Reads from the Peruvian and Mexican isolates mapped against the Chinese reference were extracted, trimmed, filtered by quality, and re-mapped, resulting in 1,317,941 reads from isolates from Puno (7,811X coverage), 5,561 from Huancayo (42X), and 674,666 from Mexico (3,395X). Genomes from Puno and Mexico were complete, while the one from Huancayo was partial. The three consensus genome sequences were of similar length (13700-13709 nucleotides). Complete mapping statistics are present in Supplementary Table 1).

Additionally, the trimmed and filtered reads were *de-novo* assembled. The same mitochondrial gene arrangement as the Chinese reference assembly was confirmed (Figure 1, Supplementary Table 2). Interestingly, the size of the protein-coding genes in Latin American samples was identical to the Chinese reference (Supplementary Table 2). An exception occurred for CYTB in the isolate from Huancayo, which has a missing codon corresponding to positions 872-874 of the Chinese reference.

**Figure 1.**
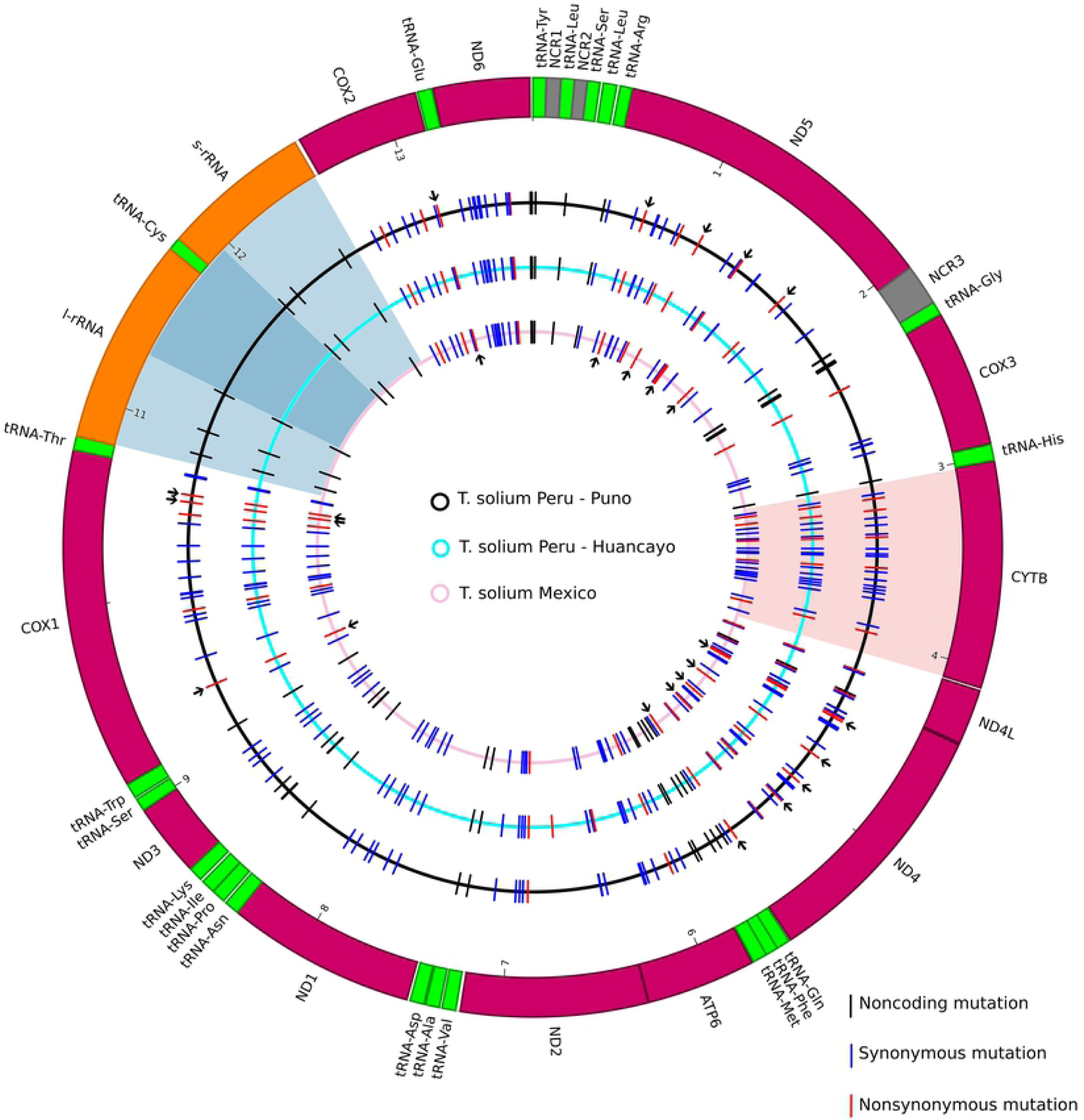
Nucleotide and amino acid differences in the 5 *T. solium genomes*. The thick outer circle is the Chinese *T. solium* reference mitochondrial genome (NCBI Reference Sequence: NC_004022.1), where inner boxes represent the coding sequences (CDS). The color code represents the CDS type: purple for protein-coding genes, green for tRNA’s, orange for ribosomal RNAs, and gray for non-coding regions. The inner circles represent the 3 *T. solium* mitogenomes assembled in this study (black: Puno, light blue: Huancayo, and pink: Mexico). The blue bars indicate synonymous nucleotide substitutions on each inner circle, while red bars highlight those causing amino acid substitutions. Flanking arrows in specific red bars indicate substitutions that involve a change in the amino acid nature. The circular segments shaded in transparent blue, and red indicate low and high variability regions, respectively. Darker blue within the blue-shaded circular segment indicates an identical region conserved in the four mitochondrial genomes. SNPs in the low variability region suggest that the region could differentiate Asian isolates from African-American isolates (as SNPs are differences with respect to the Chinese genome). However, the identical SNP distribution in all Latin American genomes suggests that it could not differentiate isolates of the same genotype, such as the African-American.

### Variability and selective pressure analysis

Variant calling, computation of the level of sequence difference (D), and selective pressure analysis were employed to provide a detailed comparison of the assembled genomes and the Chinese reference. SNP distribution detected in Latin American mitochondrial genomes was almost identical (Figure 1). Of the 34 non-synonymous SNPs, 31 were present in the three Latin American samples (Supplementary Table 3). They were distributed in all protein-coding genes except for NADH-ubiquinone oxidoreductase chain 1 (ND1) and NADH-ubiquinone oxidoreductase chain (ND3). Notably, 13 of the 31 SNPs changed the amino acid physicochemical nature. Those were located within COX1, cytochrome C oxidase subunit 2 (COX2), NADH-ubiquinone oxidoreductase chain 4 (ND4), and NADH-ubiquinone oxidoreductase chain 5 (ND5) (Figure 1, Supplementary Table 3). These change-in-nature SNPs represented 83.33% of the total mutations detected within ND5.

The similarity in the SNPs distribution along Latin American samples is also supported by the small contribution of Huancayo, Puno, and Mexico pairwise comparisons in the accumulative D (Figure 2). This was less than 0.01 for COX1, COX2, CYTB, NADH-ubiquinone oxidoreductase chain 2 (ND2), and ND5; and 0 for the other protein-coding genes (Figure 2). Five genes showed relatively high D values: ATP synthase subunit 6 (ATP6), COX2, CYTB, ND4, and NADH-ubiquinone oxidoreductase chain 6 (ND6). CYTB was the only one with nonzero values for all the pairwise comparisons.

**Figure 2.**
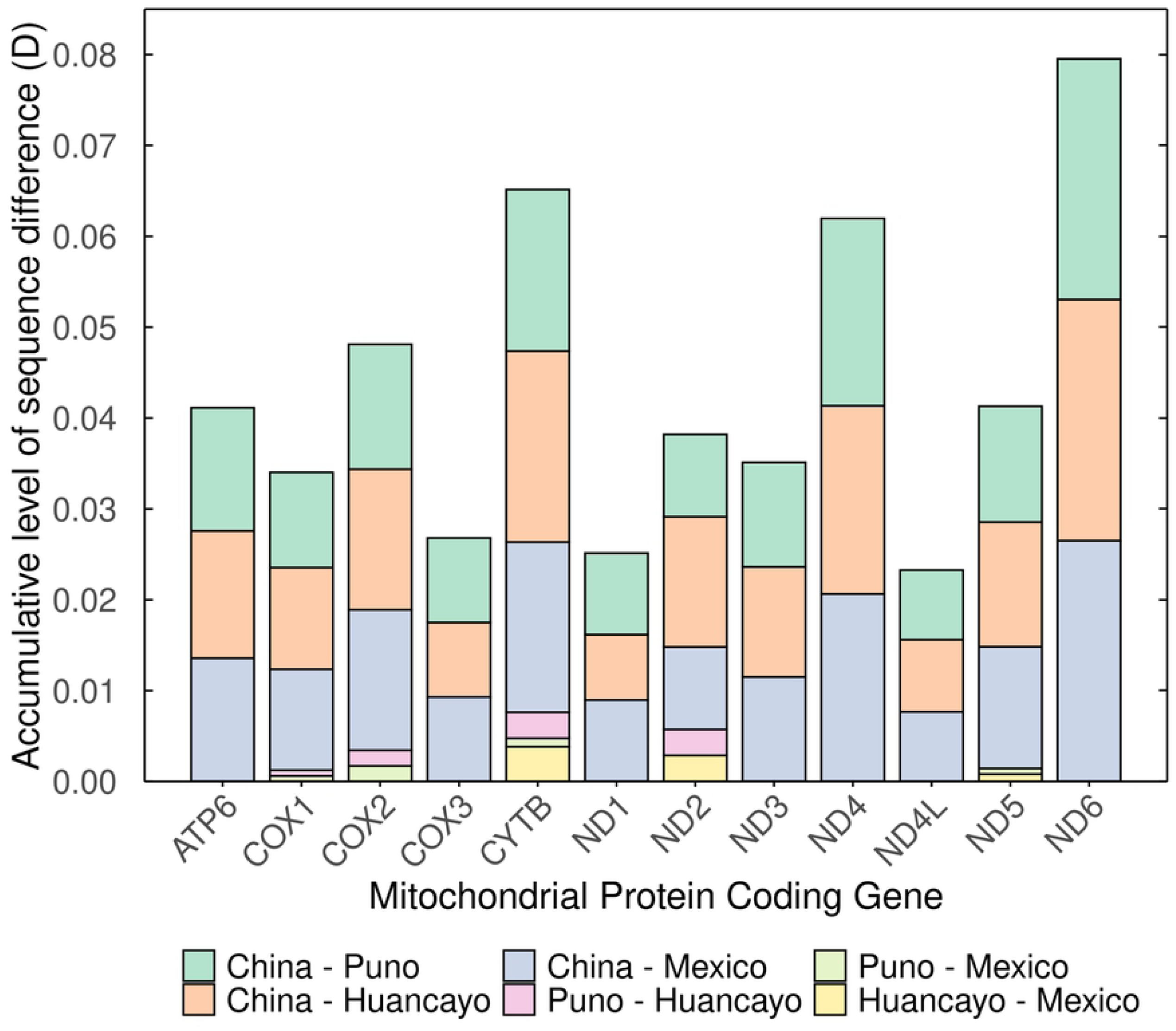
D values per protein-coding gene of each possible pairwise combination of 4 mitochondrial genomes (Chinese reference, Huancayo, Puno, and Mexico). D was used to indicate the level of sequence difference between the four mitochondrial genomes used in this study. D values were computed by performing all possible pairwise alignments and applying the D = 1 -(M/L) formula. L is the difference between the alignment length and the number of ambiguous codons. M is the number of invariant sites in the alignment. Different D values for each combination were stacked and presented per protein-coding gene in a bar plot. Thus, the height of a particular bar of a gene corresponds to the sum of D values for the different pairwise combinations; in other words, the bar height is an accumulative D value.

All the protein-coding genes presented a Ka/Ks ratio of less than 1, indicating a purifying selection. In particular, NADH-ubiquinone oxidoreductase chain 1 (ND1) and NADH-ubiquinone oxidoreductase chain (ND3) seem to be subject to absolute purification (Ka/Ks = 0) in all the samples evaluated. The three Latin American samples had the same Ka/Ks values for ATP6, cytochrome C oxidase subunit 3 (COX3), ND4, NADH-ubiquinone oxidoreductase chain 4L (ND4L), and ND6. In contrast, COX2 and ND2 had higher Ka/Ks in the samples from Puno and Huancayo, respectively, while ND5 had a higher Ka/Ks in the Mexican sample (Figure 3).

**Figure 3.**
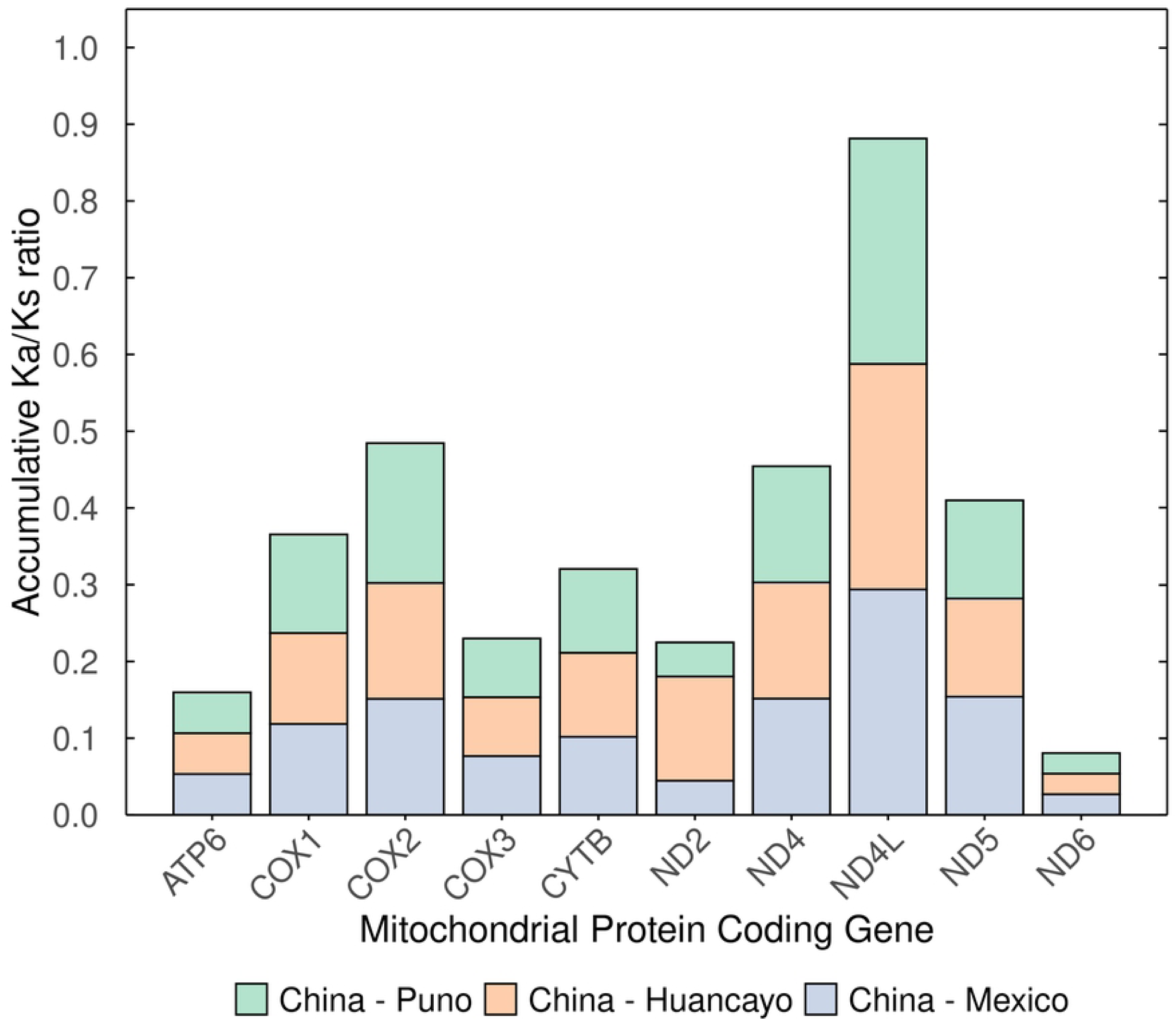
Ka/Ks ratios per protein-coding gene. Ka/Ks ratios against the Chinese reference mitochondrial genomes were calculated using the whole-genome multiple alignment in the DNAsp software for each protein-coding gene of the mitochondrial genomes. Ka/Ks values of the same protein-coding gene were stacked together (bars indicate accumulative Ka/Ks).

### Phylogenetic analysis

To identify further differentiation within the Asian and African-American genotypes, a phylogenetic reconstruction using COX1 and CYTB was performed, including the isolates of this study and others reported worldwide. The phylogenetic analysis distinguished two major clades for both markers: Asian and African-American. While both genes supported the Asian clade (COX1: PP=0.95; CYTB: BS=96%, PP=1.0), the African-American clade was only supported by the COX1 marker (BS=98%, PP=0.99). See Figure 4.

**Figure 4.**
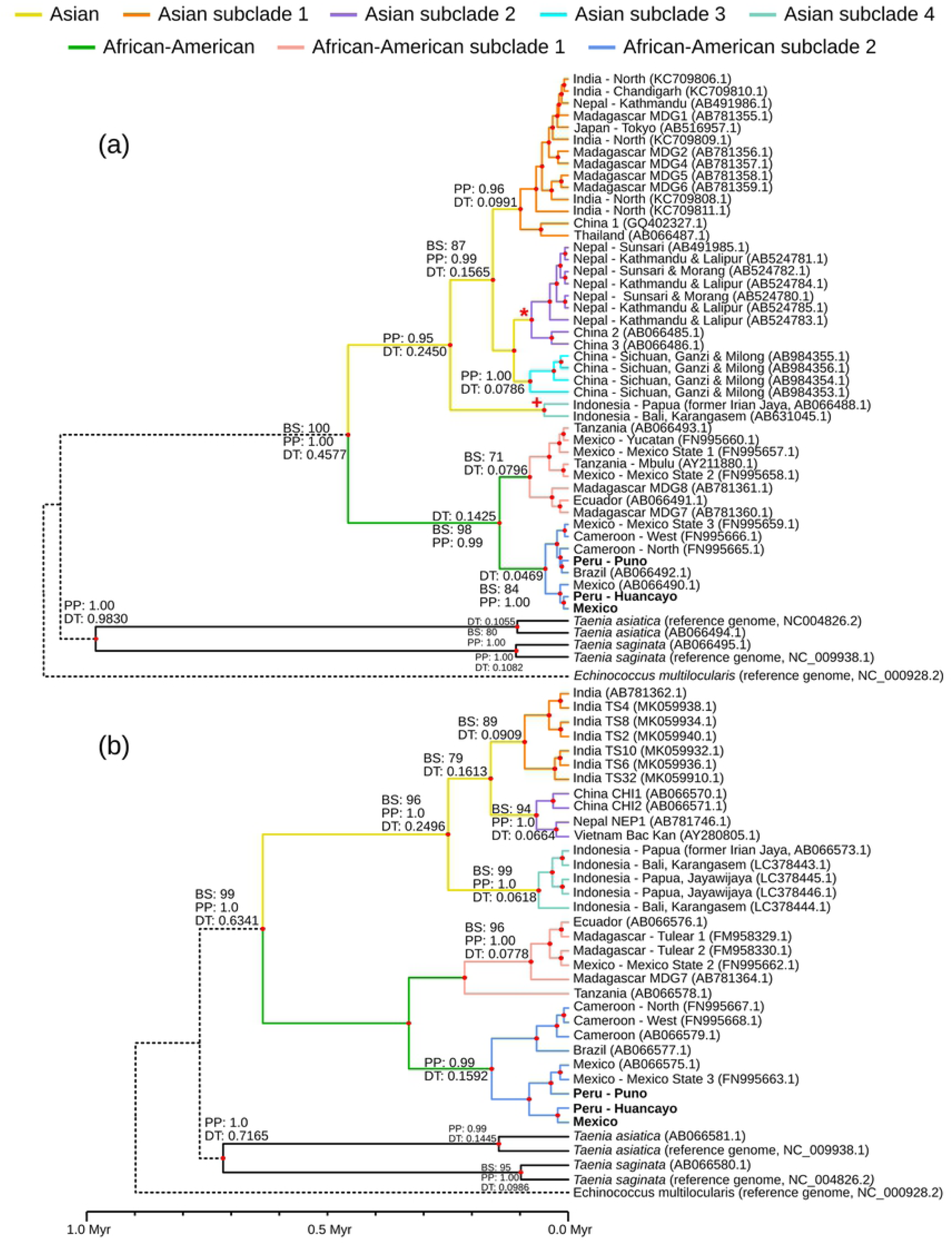
Phylogenetic trees constructed from COX1 and CYTB complete sequences. (a) Bayesian Inference (BI) tree constructed with 45 *T. solium* COX1 complete sequences. (b) BI tree constructed with 31 CYTB complete sequences. Posterior probabilities (PP) above 0.7 are shown until the level of Asian and African-American subclades. Maximum Likelihood (ML) trees were also constructed using the same sequences. If a clade has specified a Bootstrap (BS) value, it means it was present in the BI and ML tree, with a BS greater than 70% for the latter. BS values above 70% are shown only until the level of Asian and African-American subclades. For both trees, the distances of the branches are based on the timeline at a scale of one million years (Myr). Only the nodes supported in the ML and BI with BS > 70% or PP > 0.7 have their divergence times (DT) shown in Myr. Again, DT is only shown until subclades. 95% highest posterior densities are specified in Supplementary Table 4. (*) BS: 78, DT: 0.0759. (+) BS: 86, PP: 1.00, DT: 0.0495.

For COX1, the Asian group was further subdivided into four subclades with high support (Figure 4a). The first comprised countries from the East (China and Japan), South (India and Nepal), and Southeast Asia (Thailand), along with the island of Madagascar (Asian subclade 1; PP: 0.96). The second included identical sequences from Nepal and China (Asian subclade 2; BS: 78). The third comprised sequences exclusively from China (Asian subclade 3; PP: 1.0). A fourth group contained two sequences from Indonesia (Asian subclade 4; BS: 78, PP: 1.0). In the CYTB-based tree (Figure 4b), groups similar to the Asian subclades 2 (BS: 94, PP: 1.0) and 4 (BS: 99, PP: 1.0) were also present with high support. In addition, a group formed by just Indian samples was present in the CYTB phylogeny (BS: 89), which might be analogous to Asian subclade 1, as Indian sequences are only present in this subclade.

In the African-American genotype, two subclades appeared within the COX1 phylogeny. The first (African-American subclade 1; BS: 71) consisted of samples from Tanzania and Mexico (Yucatan, Mexico State 1 and 2). The second (African-American subclade 2; BS: 84 and PP: 1.0) comprised samples from Mexico (Mexico State 3 and the Mexican sequence assembled in this study), Cameroon (West and North), Peru (Puno and Huancayo), and Brazil. CYTB presented a similar topology for the African-American genotype to that obtained with COX1. The group between all the countries of the African-American subclade 1 is supported but not including Tanzania (BS = 96, PP = 0.99).

### Divergence time

Divergence times and their 95% highest posterior density intervals were calculated to situate the differentiation events within the time scale. Divergence of Asian and African-American genotypes occurred 0.458 [0.0405, 1.0625] Myra for COX1 or 0.634 [0.0667, 1.4574] Myra for CYTB.

Asian subclade 4 (Indonesia) diverged from the rest of Asia around 0.2450 [0.025, 0.5683] Myra for COX1 (Figure 4a) or 0.2496 [0.0284, 0.5804] Myra for CYTB (Figure 4b). Asian subclade 1 differentiated from Asian subclade 2 and 3 0.1565 [0.0161, 0.3526] Myra or 0.1613 [0.0200, 0.3826] Myra, for COX1 and CYTB, respectively.

The earliest divergence within Asian subclade 1 occurred at 0.0991 [0.0087, 0.2259] Myra according to COX1 or 0.0909 [0.0051, 0.229] Myra according to CYTB. For COX1, this divergence formed the common ancestor of a Chinese and Thailandese sample. Within Asian subclade 2, differentiation of Chinese samples from Nepalese (COX1) or Nepalese and Vietnamese (CYTB) occurred 0.0759 [0.0058, 0.1796] Myra for COX1 or 0.0664 [0.0035, 0.1738] Myra for CYTB. Asian subclade 3 was only present in the COX1 phylogeny. Within it, one Chinese sample diverged 0.0786 Myra.

According to COX1, the African-American subclade 1 diverged from subclade 2 about 0.1425 [0.0105, 0.3504] Myra. The clade that included both African-American subclades was not supported in the CYTB phylogeny, so their divergence time is not specified. A list of the divergence times with their 95% highest posterior density intervals are present in Supplementary Table 4.

### Haplotype network

To confirm if the samples of the subclades identified formed differentiated subpopulations, haplotype networks using COX1 and CYTB were constructed. From the COX1 multiple alignment, 43 positions of high variability were identified, supporting 25 haplotypes. These were diagrammed according to their genetic distances in a haplotype network (Figure 5). Sequences comprising each haplotype are listed in Supplementary Table 5. In contrast, the alignment of 31 CYTB sequences collapsed just into 11 haplotypes, which were generated from 23 polymorphic sites. The haplotype diversity (Hd) was 0.93 for COX1 and 0.88 for CYTB, respectively.

**Figure 5.**
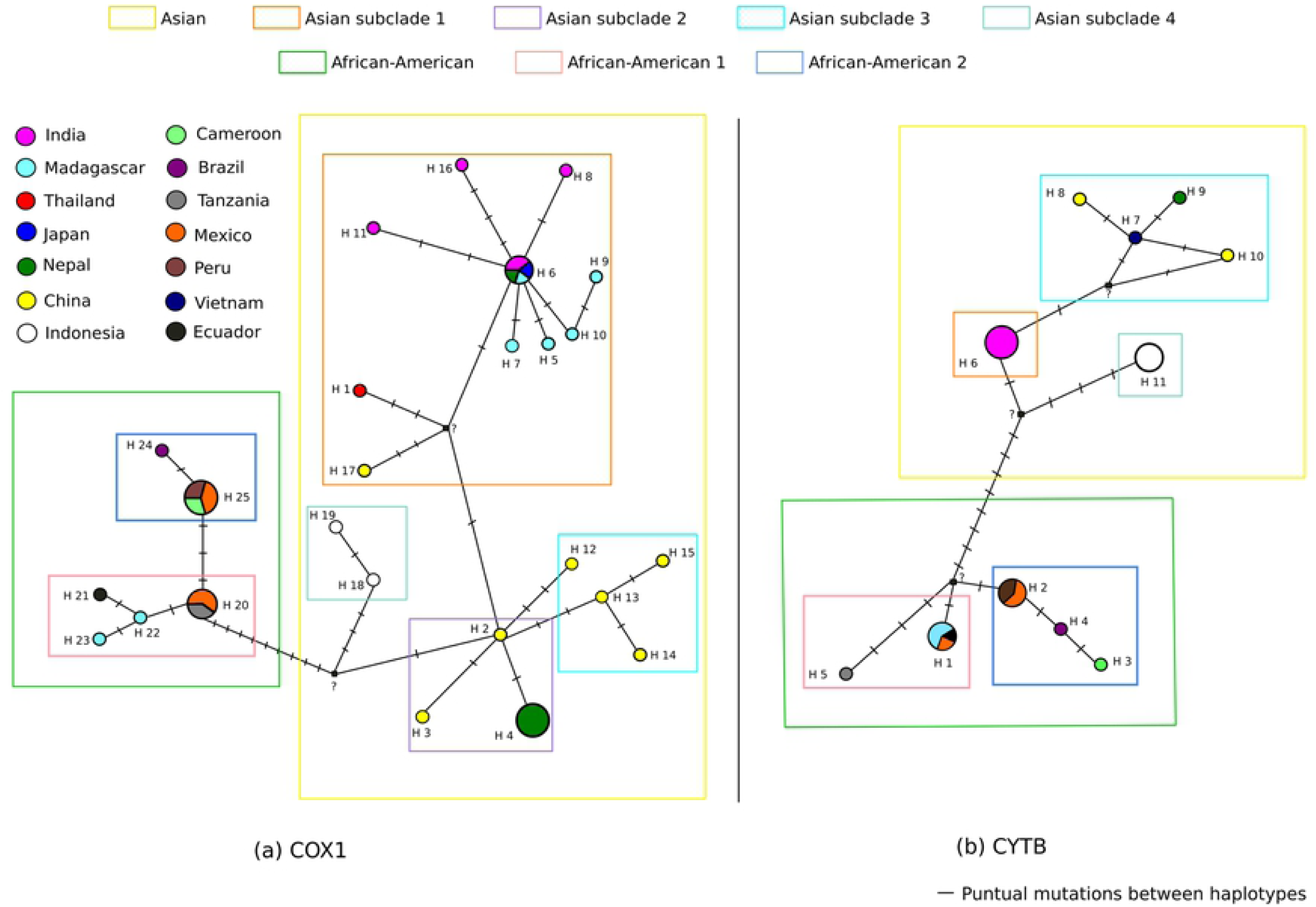
Haplotype network of COX1 and CYTB. (a) COX1 network. (b) CYTB network. The geographic origins of the samples included in each haplotype are color-coded. Colored squares indicate the most important clades and subclades.

Both haplotype networks suggested that Asian subclades 1, 2, and 3 were closely related. They formed a unique subpopulation with a dispersion center with a high Indian component. In the COX1 network (Figure 5a), haplotype 6 (H6) was the dispersion center. It was composed of sequences from Japan, India, Nepal, and Madagascar, all separated by the ocean (except Nepal and India). Around it, unique haplotypes were found distributed in India (H11, H16, H8) and Madagascar (H5, H7, H9, H10). One branch connected to an unknown haplotype, which diverged into a haplotype from Thailand (H1) and China (H17). The unknown haplotype was also connected to Chinese haplotype H2, from which other Chinese and Nepali haplotypes diverged. For CYTB (Figure 5b), the dispersion center (H6) was exclusively composed of Indian samples. H6 was connected to Chinese, Nepali, and Vietnamese samples from Asian subclade 3 through an unknown haplotype. Notably, the samples of Asian subclade 3 showed a strong interconnection between them.

Indonesian samples remained somewhat isolated from the rest of Asian countries in both networks, forming another subpopulation. For COX1, the isolate from Papua (former Irian Jaya, H18) seemed to be more basal than the one of Bali (H19). For the CYTB network, all Indonesian samples were clustered together (H11).

The COX1 network distinguished two separated groups related to the African-American subclades 1 and 2, which differentiated in 3 punctual mutations. The CYTB network also distinguished between African-American subclades. However, it separated a haplotype conformed by a unique Tanzanian sample from the rest of the countries of African-Subclade 1. To determine if the countries that formed African-American subclade 1 were genetically different from those of the African-American subclade 2, a computation of the ϕst value between these two groups was made. Values were 0.83 for COX1 and 0.62 (p < 0.05) for CYTB.

## DISCUSSION

The mitochondrial genomes from Puno and Mexico had 7,811X and 3,395X of coverage, respectively. They had no ambiguous nucleotides and were the same size as the reference. The sample from Huancayo had a lower coverage (42X). Although this resulted in a partial genome, it was still adequate for the rest of the analysis. As expected, the genome size and the %GC are similar between these three isolates, supporting the assembly method. No gene rearrangements were detected (Figure 1, Supplementary Table 2), and the size of each protein-coding gene is the same, except for the gene CYTB in Huancayo, which has one codon less (Supplementary Table 2). Nakao *et al*. [18] previously reported the presence of an abbreviated stop codon U (or T in DNA) at the ND1 gene. Noteworthy, all the mitochondrial genomes assembled in the present study present this stop coding, confirming this observation.

Variant calling showed that differences with respect to the Chinese reference are almost identical in all the Latin American genomes and concentrated in protein-coding genes, as revealed by the SNPs distribution (Figure 1). For instance, of the 34 non-synonymous SNPs, 31 are present in all the assembled genomes (Supplementary Tables 3).

Surprisingly, almost half of the Latin American SNPs involve a change in the amino acid physicochemical nature (Figure 1, Supplementary Table 3). Those are located in four mitochondrial protein-coding genes and especially in ND5. It is well-established that change-in-nature mutations affect the structure and function of proteins [33]. Hence, the change-in-nature SNPs in the Latin American samples could have altered the folding of their mitochondrial proteins compared to the proteins of Asian *T. solium*. This may be linked to a reduced aerobic respiration efficiency that may affect their fitness in oxygen-rich environments. The situation above-described could lead to a negative tropism towards the oxygen-rich subcutaneous tissue, which has an O_2_ partial pressure of 40-80 mmHg [34] higher than the ∼23 mmHg [35] and <11 mmHg [36,37] of the brain and intestinal lumen, respectively. Accordingly, it has been suggested that subcutaneous cysticercosis is uncommon in Latin America but not in Asia [38–40]. The evidence reported here calls for studies to confirm this clinical difference between regions and relate it to the change-in-nature SNPs found.

The present results suggest that the CYTB sequence is the most variable of the whole mitochondrial genome. It has the highest density of SNPs, the highest accumulative D, and different D values for all pairwise comparisons (Figure 2). Besides, it has a medium length of 1068 bp (Supplementary Table 2). Taking these features together show that CYTB is a more suitable molecular marker for intraspecific variability analysis that will differentiate isolates of the same or different genotypes. In agreement, it has been stated that, within *taeniidae*, CYTB is a better marker for reconstructing phylogenies among closely related groups (such as intraspecific variations) because of its higher evolutionary rate [15]. Indeed, other studies have reported results that support that CYTB has higher variability than, for example, COX1. For instance, 28 SNPs (1.7% variability rate) in the COX1 gene were found in contrast to the 31 SNPs (2.9% variability rate) present in CYTB [6]. Despite this, more complete sequences for COX1 are available compared to CYTB. Additionally, COX1 sequences have a greater variety of geographical origins. Extra CYTB sequences would allow better and more informative phylogeographic reconstructions of *T. solium* lineages.

ATP6, COX2, and ND6 also have a relatively high D value; however, the three of them have a short length. D values could overestimate the variability for small genes as the percentage of identity is inversely correlated to the alignment length. Thus, high D values for small sequences as these three should be taken cautiously and do not necessarily imply high variability.

The region corresponding to the small and large ribosomal RNA (rRNA) and the cysteine transfer RNA (tRNA-Cys) showed the lowest quantity of SNPs. This and the fact that these SNPs are present in all the Latin American genomes suggest the low variability of the region. Moreover, this region contains an internal sequence that remains identical among the Chinese and Latin American genomes. The internal sequence could be the target of conserved primers to specifically amplify the *T. solium* mitochondrial DNA, as some portions differ from *T. saginata* and *T. asiatica* (Supplementary Figure 1).

ND1 and ND3 are subjected to absolute purification (Figure 3), which is corroborated by the fact that only synonymous SNPs were detected. In that sense, mutations in these genes seem deleterious and therefore negatively selected. ND1 is a crucial subunit of the mitochondrial respiratory complex I because it allocates the entrance of the quinone reaction chamber and the first half part of the first proton translocation channel, which receives input from the cytoplasm [41,42]. Moreover, ND3 provides the flexibility needed for a concerted rearrangement that generates the driving force for proton pumping [43]. Hence, mutations in these genes could affect the quinone reductase activity and collapse the proton translocation system on the inner mitochondrial membrane. The importance of both subunits is in agreement with the fact that these genes are “cold spots” for amino acid mutations.

The phylogenetic analysis of both genes showed two main lineages: the Asian and the African-American (Figure 4), which has been reported by other studies [6,8,13]. Divergence between those lineages occurred 0.458 [0.0405, 1.0625] (COX1, Figure 4a) to 0.634 [0.0667, 1.4574] (CYTB, Figure 4b) Myra during the Pleistocene, in agreement with previous works [6,9,13].

One study has reported archaeological evidence that places modern humans in Daoxian, China, 0.12 to 0.08 Myra [44]. However, it has not been confirmed if this was a successful (persistent) settlement in south Asia. Interestingly, the present results suggest that Chinese *T. solium* started to diverge around a similar period, 0.0786 to 0.0664 Myra according to Asian subclades 2 (both phylogenies, Figure 4a,b) and 3 (COX1 phylogeny, Figure 4a). The divergence within a geographical region requires this region to be persistently settled by humans infected with *T. solium*. Hence, the present data suggest that humans successfully populated south China around this period and that the archaeological evidence might have arisen from these early settlements.

The phylogenetic analyses suggested that the Asian genotype is further subdivided into four (COX1) or three (CYTB) subclades. However, both haplotype networks (Figure 5) indicated that Asian subclades 1, 2, and 3 are closely related, forming a unique subpopulation with India as the center of dispersal [7]. That subpopulation is differentiated from another formed by Indonesian samples. The different degrees of relationship between Asian *T. solium* samples suggest heterogeneous gene flow.

In the COX1 haplotype network (Figure 5a), samples from Japan, India, Nepal, and Madagascar are grouped in the same haplotype (H6). Considering that *T. solium* is not endemic in Japan, its relation with H6 samples is likely the result of a recent reintroduction, a not-so-rare event in the last years [45]. The Madagascan isolates’ close association with Nepali and Indian samples suggests that the parasite was introduced into Madagascar from the Indian subcontinent [12]. A particular case occurred with Nepali samples. One group of samples was included in H6 in close association with Indian isolates; however, the other was included in H4 in close association with Chinese sequences. These two genetic subpopulations suggest the existence of two gene flows towards Nepal, one from the north (from China) and another from the south (from India). They remain separated, possibly due to the geographical barrier that the Himalayas constitute.

Asian subclade 4, which is formed by Indonesian samples, was the first to diverge ∼0.25 Myra according to COX1 and CYTB phylogenies. A similar divergence time was found in previous research [9]. This correlated with a basal position concerning the other Asian subclades in both phylogenies, which has also been reported [6,9]. Interestingly, both haplotypes networks grouped Indonesian samples in haplotypes that remain distant from other Asian sequences. Furthermore, both haplotype networks directly linked them to an unknown haplotype, which bridged the African to the Asian lineages. Given that the unknown haplotype is the only bridge between the two genotypes, it is proposed that it corresponds (or is related) to the *T. solium* modern ancestor that independently dispersed in Asia and Africa. These results suggest that Indonesian samples are more ancient and related to the modern *T. solium* ancestor than other Asian samples. This could be explained by the introduction of the parasite by early human migrations followed by isolation of the population (lack of gene flow). Studying more *T. solium* samples from Indonesia could help identify plesiomorphies and, by comparing them with more recent isolates, obtain an insight into how *T. solium* evolves.

Indeed, *T. solium* subpopulations in Indonesia are isolated, as shown by the fact that this parasite is restricted to Bali and Papua (former Irian Jaya). The COX1 haplotype network suggested that the sample from Papua is more ancient than the one of Bali because it is closer to other Asian isolates. Nonetheless, this proposal seems inconsistent with the epidemiological evidence that suggests that the introduction of *T. solium* to Papua was made 50 years ago from Bali [14,46]. Interestingly, even though CYTB is more variable than COX1, the CYTB network shows no resolution to distinguish between Indonesian haplotypes, while the COX1 network does. This inconsistency might suggest that the haplotype differentiation seen in the COX1 haplotype network was an artifact attributed to a random selection when *T. solium* was introduced into Papua, as it has been hypothesized in other work [14].

Remarkably, phylogenies and haplotype networks constructed in this work suggest that the African-American lineage is further subdivided into two groups (African-American subclades 1 and 2). The genetic differentiation between the two is confirmed by the fact that ϕst values were high and significant (p < 0.05). Interestingly, the subdivision was observed in a previous study [8]. Nonetheless, they did not include African samples, making it impossible to perform inferences about the origin of both subclades. The haplotype networks further confirmed the subdivision, allowing one to visualize two different clusters.

Regarding African-American subclade 2, both haplotype networks showed a close relation between samples from Brazil, Peru, Mexico, and Cameroon, considering the majority were included in the same haplotype. Of note, this group only had sequences from West Africa (Cameroon). The geographic composition and degree of association of the samples of this subclade are coherent with the conversion between the Trans-Atlantic slave and trade routes [7]. These trade routes mainly connected West Africa, Europe, and the Americas.

As for the African-American subclade 1, the COX1 network revealed a close association between samples from Mexico, Tanzania, Madagascar, and Ecuador. The low differentiation and high genetic flow between one Mexican and one Tanzanian isolate were reported previously [7]. Of note, this group exclusively included sequences from East Africa (Tanzania and Madagascar). East African countries were not direct participants in the Trans-Atlantic trade routes. Hence, their link with Latin American samples might have been caused by another source of gene flow, such as the one generated by the Indian Ocean slave trade. This trade connected East Africa, Indian Ocean countries, the Middle East, and later the Americas.

All in all, the subdivision of the African-American genotype reveals that two different sublineages exist in Latin America: one that derives from West Africa and the other originated in East Africa. Two separate gene flows created from the Trans-Atlantic and Indian Ocean trade routes may have caused the observed distribution. Notably, Mexican samples were present in both lineages, which agrees with the proposal that at least two genetic subpopulations coexist in Mexico [13]. Although the Peruvian isolates were only included in the East African linage, subpopulations from the West African linage are not discarded to exist, given that Mexico and Peru were trade centers during European colonization.

In conclusion, thirteen SNPs with respect to the Chinese reference that involved a change in the amino acid physicochemical nature were identified. They might be related to differences in aerobic respiration efficiency between Asian and Latin American *T. solium*. Further differences within the Asian genotype were also reported. For instance, its differentiation and basal position combined with its divergence time suggest that the subclade formed by Indonesian samples is the most ancient and closely related to the modern *T. solium* ancestor than other Asian sequences. Strikingly, all phylogeographic analyses showed that the African-American genotype is subdivided into two subgroups. One has a strong relation with East African countries while the other with West Africa, which might reflect the influence of the Trans-Atlantic and the Indian Ocean trade routes. Of note, the isolates whose genomes were assembled were part of the West African sublineage. In summary, the present study shows that a detailed comparison of the mitochondrial variability of *T. solium* within and between Asian and African-American genotypes still reveals interesting features that could be used to combat *T. solium* diseases.

## DATA AVAILABILITY

The assembled and annotated mitochondrial genomes from Puno and Huancayo were uploaded to the GeneBank with accession numbers IN_PROCESS_OF_SUBMISSION and KT591612, respectively. Regarding the assembled genome from Mexico, nucleotide sequence data reported are available in the Third Party Annotation Section of the DDBJ/ENA/GenBank databases under the accession number TPA: BK061219. Other raw data will be available upon request.

## ACKNOWLEDGEMENTS

We acknowledge Luis Tataje, Manuel Ramirez, and Hugo Valdivia for their advice during the genome analysis. We are also grateful to Lorraine Michelet for her invaluable assistance and share of her doctoral thesis that helped complement the conception of this research.

## SUPPLEMENTARY MATERIAL LEGENDS

**Supplementary Figure 1. Selected portions of the multiple alignment of the identical internal region**. The sequences corresponding to the identical internal region conserved in the *T. solium* mitochondrial genome from China (NC_004022.1), Puno, Huancayo, and Mexico (see Figure 1) were aligned with the mitochondrial genomes of *T. asiatica* (NC_004826.2), *T. saginata* (NC_009938.1) and *E. multilocularis* (NC_000928.2) in the online version of Clustal Omega [23]. The alignment was then cropped to include only matching sections using CLC Genomics Workbench v. 21.05.5 (https://digitalinsights.qiagen.com/). Selected portions of the alignment that showed significant differences between *T. solium* and the other organisms are presented.

**Supplementary Table 1. Mapping statistics of the genome assembly**. The number of reads mapped to the Chinese mitochondrial genome (reference) and their mean quality scores before trimming (pre) are given. The number of reads mapped after trimming (post), the coverage, the genome length in base pairs (bp), the %GC, and the N50 of the assembly in nucleotides (nt) are also shown.

**Supplementary Table 2. Gene arrangement of *T. solium* mitochondrial genomes from Peru, Mexico, and China (reference)**. The size of each genome in base pairs (bp) is given in parenthesis next to the mitochondrial genome name. Position intervals per gene are shown. The size of each gene (in bp) is specified in parentheses next to each position interval. Start and stop codons of each protein-coding gene per mitochondrial genome are also specified.

**Supplementary Table 3. Non-synonymous mutations in the Latin American mitochondrial genomes with respect to the Chinese reference**. Amino acid mutations in all the assembled genomes of this work are given per protein-coding genes. In the present work, standard amino acids were classified into five groups, considering their major species at pH of 7: polar and uncharged (S, T, N, and Q), polar and positively charged (R, H, and K), Polar and negatively charged (D, E), nonpolar (A, V, I, L, M, F, Y, and W) and special cases (C, G, and P). A mutation that changes the amino acid nature involves that the mutant amino acid belongs to a different group than the original. Mutations highlighted in red and blue represent changes that do and do not involve a change in the amino acid nature, respectively.

**Supplementary Table 4. Divergence events and their dating**. Divergence events in COX1 and CYTB phylogenies are described. The gene in whose phylogeny the event occurred is specified in bold at the start of the divergence event description. If nothing is specified, the same event occurred in both phylogenies. The divergence times with 95% highest posterior density intervals (brackets) in millions of years ago (Myra) are also reported for each event.

**Supplementary Table 5. Sequence composition of the haplotypes formed in the COX1 and CYTB networks**. Sequences (with their accession numbers) included in each haplotype are specified.

